# A Technical Evaluation of Plasma Proteomics Technologies

**DOI:** 10.1101/2025.01.08.632035

**Authors:** William F. Beimers, Katherine A. Overmyer, Pavel Sinitcyn, Noah M. Lancaster, Scott T. Quarmby, Joshua J. Coon

## Abstract

Plasma proteomic technologies are rapidly evolving and of critical importance to the field of biomedical research. Here we report a technical evaluation of six notable plasma proteomic technologies – unenriched (Neat), Acid depletion, PreOmics ENRICHplus, Mag-Net, Seer Proteograph XT, Olink Explore HT. The methods were compared on proteomic depth, reproducibility, linearity, tolerance to lipid interference, and limit of detection/quantification. In total we performed 618 LC-MS/MS experiments and 93 Olink Explore HT assays. The Seer method achieved the greatest proteomic depth (∼4,500), while Olink detected ∼2,600 proteins. Other MS-based methods ranged from ∼500-2,200. Neat, Mag-Net, Seer, and Olink had strong reproducibility, while PreOmics and Acid showed higher variability. All MS methods showed good linearity with spiked-in C-Reactive Protein (CRP); CRP was surprisingly not in the Olink assay. None of the methods were affected by lipid interference. Seer had more than double the number of quantifiable proteins (4,800) for both LOD and LOQ than the next best method. Olink was comparable to Neat and Mag-Net for LOD, but worse for LOQ. Finally, we tested the applicability of these methods for detecting differences between healthy and cancer groups in a non-small cell lung cancer (NSCLC) cohort.

## Introduction

Owing to its ease of collection, organism wide circulation, and rich molecular content, blood offers an attractive medium by which to monitor the physiological state. Human blood, for example, contains myriad species including metabolites, lipids, nucleic acids, and proteins – many of which are currently used as clinical markers of health^1–4^. Thousands of proteins, the ultimate functional drivers for most biological processes, can be found in blood plasma^5,6^. Many of the proteins have their primary function in the circulatory system, while thousands of other non-secreted proteins may result from infection, disease, and tissue leakage^7^. Yet characterizing and measuring the plasma proteome has been complicated by the extremely broad dynamic range over which these proteins occur (∼ 10^12^). This challenge has both limited the scope of plasma proteome discovery and spurred technological innovation.

Mass spectrometry methods currently offer the most in-depth and global proteome analyses^8–10^. However, the dynamic range of the plasma proteome has historically limited the achievable depth to a few hundred proteins as compared to thousands when analyzing cells or tissues^11^. MS advancements including the development of data-independent acquisition (DIA)^12–14^ alongside fast and high-resolution mass analyzers have boosted the achievable plasma proteome depth by a factor of two to three-fold^15,16^. While these innovations have been impactful, alone they still only allow detection of a small fraction of the plasma proteome. Concomitant to the MS innovations, sample preparation developments have similarly accelerated. Initial efforts focused on the use of antibodies to deplete high abundance proteins but they only marginally improved depth and suffered from irreproducibility^17–19^. In 2020, Blume et al. first described the use of functionalized nanoparticles to form protein coronas and thereby capture an enriched plasma proteome for downstream MS analysis^20^. This technology boosted the plasma proteome coverage by five-fold to ∼ 2,000 proteins. That technology has been further refined and is commercially available by Seer (Proteograph XT). In 2023, two other methods involving magnetic particle-based enrichment for plasma proteomics were described – ENRICH technology by PreOmics and Mag-Net by the MacCoss laboratory and ReSyn Biosciences^21^. While Mag-Net aims to enrich extracellular vesicles, ENRICH technology targets low abundance proteins. Also in 2023, Steen and colleagues described use of perchloric acid to generate a depleted supernatant for MS analysis^22,23^. Note, this is not a comprehensive list of all the MS-based plasma proteomic technologies but rather represents notable recent advances.

Affinity-based approaches have long been used for low-throughput plasma protein detection. Over the past decade, however, two companies, Olink and SomaLogic, have pioneered affinity-based global plasma protein detection^24–29^. These technologies leverage libraries of affinity reagents to target specific proteins. Olink, for example, uses antibody pairs for each protein target (i.e., proximity extension assay, PEA). When the antibody pairs bind in close proximity, the attached DNA ligates and is amplified and analyzed using next-generation sequencing^26^. Presently, Olink targets 5,415 proteins in their Explore HT assay.

Given the potential of these technologies to detect and quantify thousands of previously inaccessible plasma proteins, providing biological insight and potential therapeutic targets, we sought to directly compare these methods. Several studies have offered interesting but limited cross-method comparisons^30–32^. Given the critical need for future plasma proteome studies to understand the pros and cons of the various approaches, we aim to provide here a rigorous comparison of six plasma proteomic technologies – Olink Explore HT and five MS-based methods. The MS methods are neat (control or non-enriched plasma preparation)^33^, Acid (perchloric acid method), Mag-Net, PreOmics ENRICHplus (pre-released kit), and Seer Proteograph XT.

To compare these six plasma proteomics methods we devised five distinct experiments, each aimed at probing a different aspect of method performance. First, we assessed proteomic depth and reproducibility using five preparation replicates. Second, to examine quantitative linearity we spiked-in an endogenous protein at various levels. Third, we tested for tolerance to lipid interference by spiking in triglycerides. Fourth, we globally assessed quantitative performance by generating a matrix-matched calibration curve. Finally, we benchmarked method performance with a 40-sample study of non-small cell lung cancer (NSCLC). Altogether, we conducted 618 LC-MS/MS experiments and an additional 93 Olink Explore HT analyses. From this data we generated a comprehensive assessment and comparison of all six plasma proteomic methods.

## Results and Discussion

To compare the six plasma proteomic methods – Neat, Acid, PreOmics, Mag-Net, Seer, and Olink – we used the same starting plasma sample split into aliquots. For the MS-based methods we utilized the same experimental setup described below, except where noted. Briefly, we used an identical LC-MS/MS method consisting of an in-house capillary nanoLC column with integrated emitter tip coupled with a nUHPLC and Orbitrap Astral hybrid MS system^34^. The Orbitrap Astral was operated in DIA mode, with each nLC-MS/MS experiment taking just under one hour from sample to sample (39-minute separation)^8,10^. Representative chromatograms from the various MS-based plasma preparation technologies are shown in **Supplemental Figure 1**. We also note that the Seer Proteograph XT protocol uses two distinct nanoparticle sets (NPA and NPB) resulting in two separate sample injections. In our hands the results from NPA alone were comparable to the aggregate of the two (**Supplemental Figure 2**); however, for all experiments in this study we combined NPA and NPB data so each Seer result represents the sum of these two one-hour experiments. The other MS-based methods (i.e., Neat, Acid, PreOmics, and Mag-Net) were all performed with single one-hour analyses. All raw MS files were analyzed using Spectronaut 19 in direct DIA mode with default settings and filtered to a 0.01 protein group q-value (i.e., 1% protein FDR)^35^. For the Olink we selected the Explore HT method (comprising 5,415 targets) and sent the samples to the contract research organization Life and Brain GmbH. Raw data received was then processed using the OlinkAnalyze R package whereby proteins below Olink suggested LOD were considered missing values. Specifically, Olink includes negative controls for every protein assay, where the PEA reagents are placed in buffer to monitor any background noise, allowing for an assay specific LOD calculation.

### Plasma proteomic depth

To evaluate the plasma proteomic technologies we first aimed to test proteomic depth and reproducibility. To accomplish this, we prepared five technical replicates of the same plasma sample (BioIVT) using each method. **Figure 1a** presents the number of identified protein groups across the methods. From this data we see a broad range of performance from ∼ 500 to 4,600 detected proteins. To assess reproducibility of identifications we include the number of proteins detected in all five replicates, the average number of proteins identified per replicate, and the total number of proteins identified across all replicates. Note the reproducibility of all the methods is high, with Olink having the largest variation in identification count. We observe that Seer detects ∼ 4,500 proteins while all the other methods achieve considerably less depth. 6,393 protein groups were identified across all methods. Next, we sought to explore the complementarity of the Olink and MS-based methods (**Figure 1b**). To avoid complications that can arise during protein grouping, we mapped each UniProt ID to its corresponding gene name and compared the overlap of all the resulting gene names. Here we see that 5,161 gene products were observed via nLC-MS/MS methods and 2,644 with Olink, but only 1,412 of these gene products were observed in both, less than 25% of the total. This proportion is almost identical to the overlap between Seer and Olink, the two largest groups, indicating that the proteins identified with Seer encompass most of the MS-based method identifications. **Supplemental Figure 3a** further explores the uniqueness of protein identifications from each of the methods.

**Figure 1.**
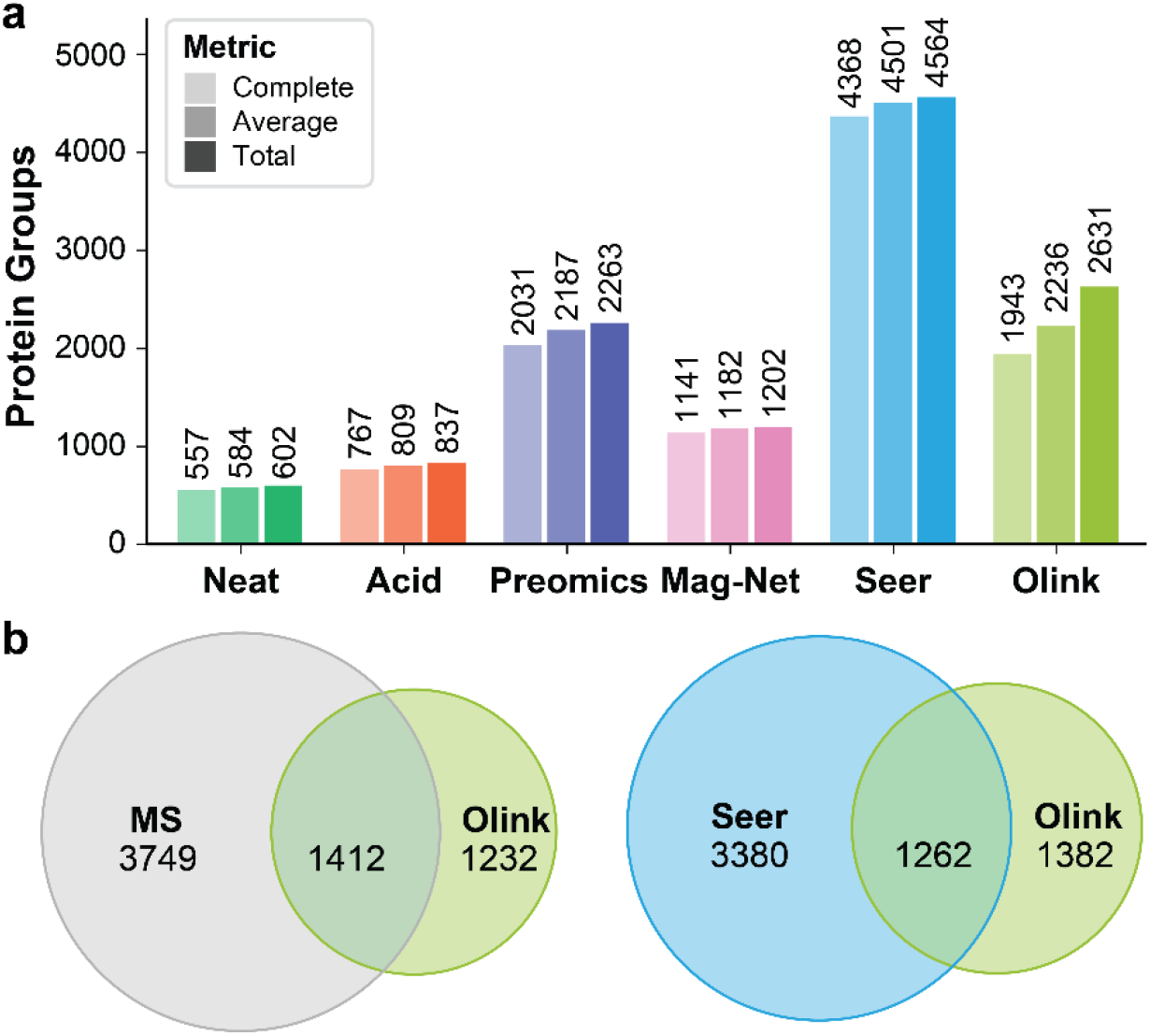
Depth of proteomic analysis afforded by the tested methods. (a) Number of protein groups identified across five technical replicates. For Olink, protein IDs were reported for assays that were above LOD. Complete identifications represent the number of protein groups detected in all five replicates, average was the average number of protein groups detected per replicate, and total were all unique protein groups detected in at least one replicate. (b) Venn diagrams showing the gene-level overlap in identifications between Olink and proteins identified in at least one of the MS methods or Olink and Seer.

To understand more about the detected proteins we utilized the Human Plasma Proteome Project (HPPP) database^6^. The HPPP has a dataset that contains 5,092 canonical plasma proteins and estimates of their abundances. Not all the 6,393 proteins detected here were contained in the HPPP dataset. We also note that the HPPP dataset was built almost entirely from nLC-MS/MS data, so it is not surprising that there is much more concordance with the MS-based methods (**Supplemental Figure 3b**). **Figure 2a** presents the overlap with the HPPP plasma proteome compendium for each of the methods. Each point represents a detected protein and the location on the x-axis shows its HPPP abundance rank. **Figure 2b** shows the median detected HPPP protein abundance across the methods. Interestingly, there were many extremely high abundance proteins that were not included in the assay list of Olink Explore HT, in part skewing the median for Olink to the lowest abundance of the methods tested.

**Figure 2.**
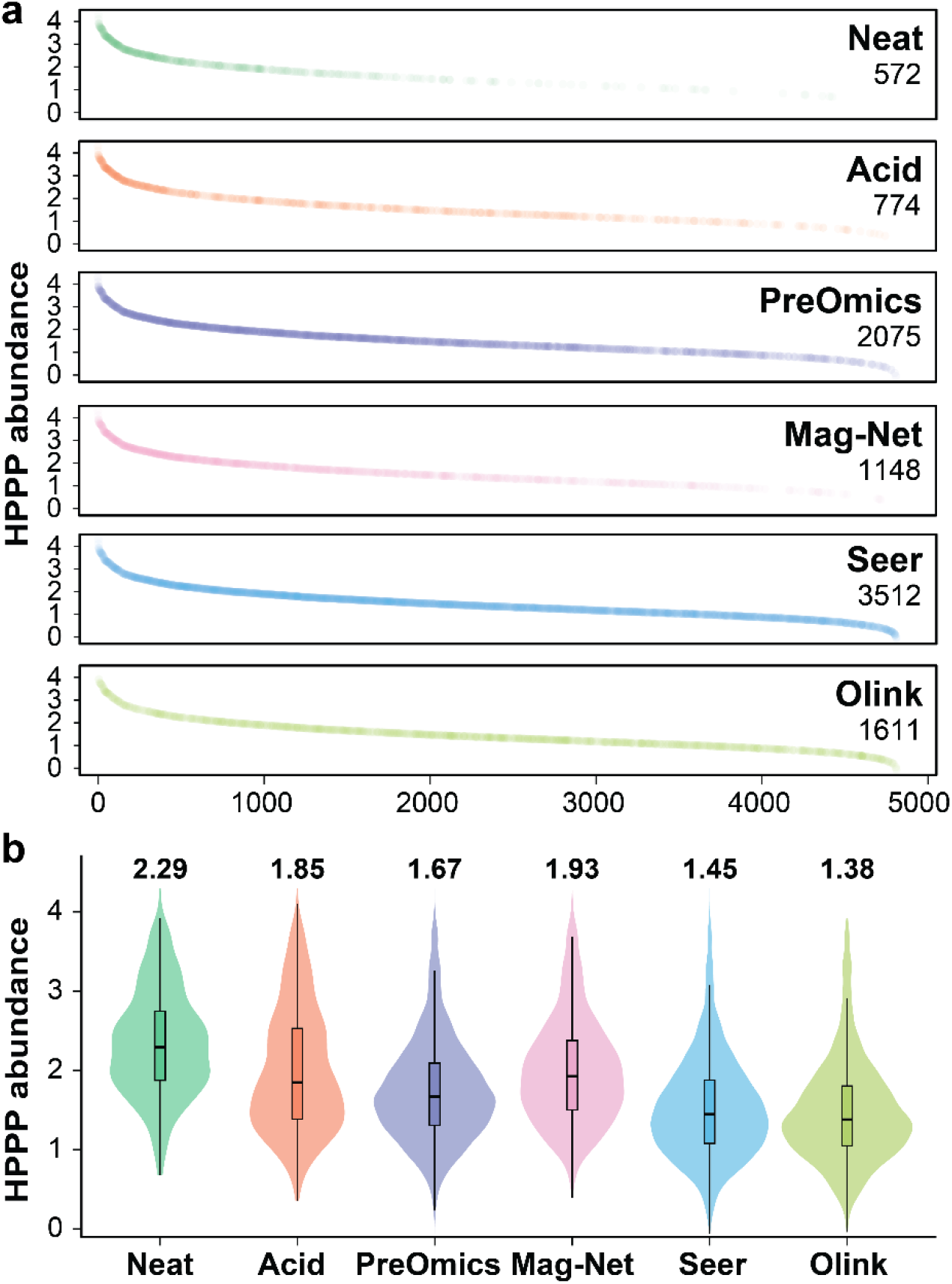
Distribution of identified proteins by Human Plasma Proteome Project estimated abundances. Proteins that were identified in common between the methods in this study and the Human Plasma Proteome Project database (n = 5091) were mapped to HPPP abundance data. HPPP abundance is calculated from log10 transformed PSM spectral counting of each protein. (a) Ranking plots with HPPP abundance on the y-axis and rank on x-axis, revealing that some methods skew toward detection of high-abundance proteins while others are more evenly distributed across abundance range. (b) Violin plots showing the density of where identifications fall in the abundance level by method.

For the MS-based methods we wondered whether the various protocols might affect the quality of the resulting peptides given that plasma has many endogenous proteases. To investigate, we examined both the trypsin missed cleavage rate and non-specific cleavage rate. We found that only about 12 to 16% of the PreOmics, Seer, and Acid prepared peptides resulted from missed tryptic cleavages while that number increased to 23 and 25% with the Mag-Net and Neat methods, respectively (**Supplemental Figure 4a**). For all methods except Acid, ∼1% of peptides resulted from non-specific cleavages while that number rose to 7% for the Acid approach (**Supplemental Figure 4b**).

### Quantitative reproducibility

Having established baseline performance for achievable proteomic depth, we turned our attention to assessing the reproducibility of each method. Using the same data presented above, we evaluated the median coefficient of variation (CV) values for each measured protein using LFQ values (MS methods) or NPX (Normalized Protein eXpression) (Olink). From **Figure 3A** we observe a wide range of performance in terms of reproducibility. The strongest performances are from the Neat (8.69%) and Seer (10.38%) methodologies, followed closely by Mag-Net (12.55%) and Olink (13.9%). PreOmics and Acid methods had high median protein CV values at 25.24% and 26.55%, respectively. In general, one would expect that the higher abundance proteins would be more reproducibly measured, as there is a higher signal/noise ratio for more reliable quantification. To investigate, and to be sure that the methods that obtain deeper coverage are not unfairly punished by the calculation described above, we next compared the median CV values for only the protein groups that were detected across all the methods (n = 137, **Figure 3b**). For most methods we see a sharp decrease in median CV value, as expected; however, the PreOmics data shows an increase in median CV values up to 48.5% while the Acid median CV remained unchanged. To more closely evaluate the relationship between protein abundance and measurement reproducibility we next plotted protein abundance vs. CV (**Figure 3c**). The shared proteins are indicated in blue, whilst all others are black. In general, we see that as the protein abundance rises, the quality of the measurement improves, except for a subset in the Acid and PreOmics data. Note, Olink uses a different quantitative metric, Normalized Protein Expression (NPX) that modifies sequencing counts with several correction factors. The CV values for Olink had no apparent correlation with NPX alone, so the proteins for all methods were again mapped to the HPPP abundances.

**Figure 3.**
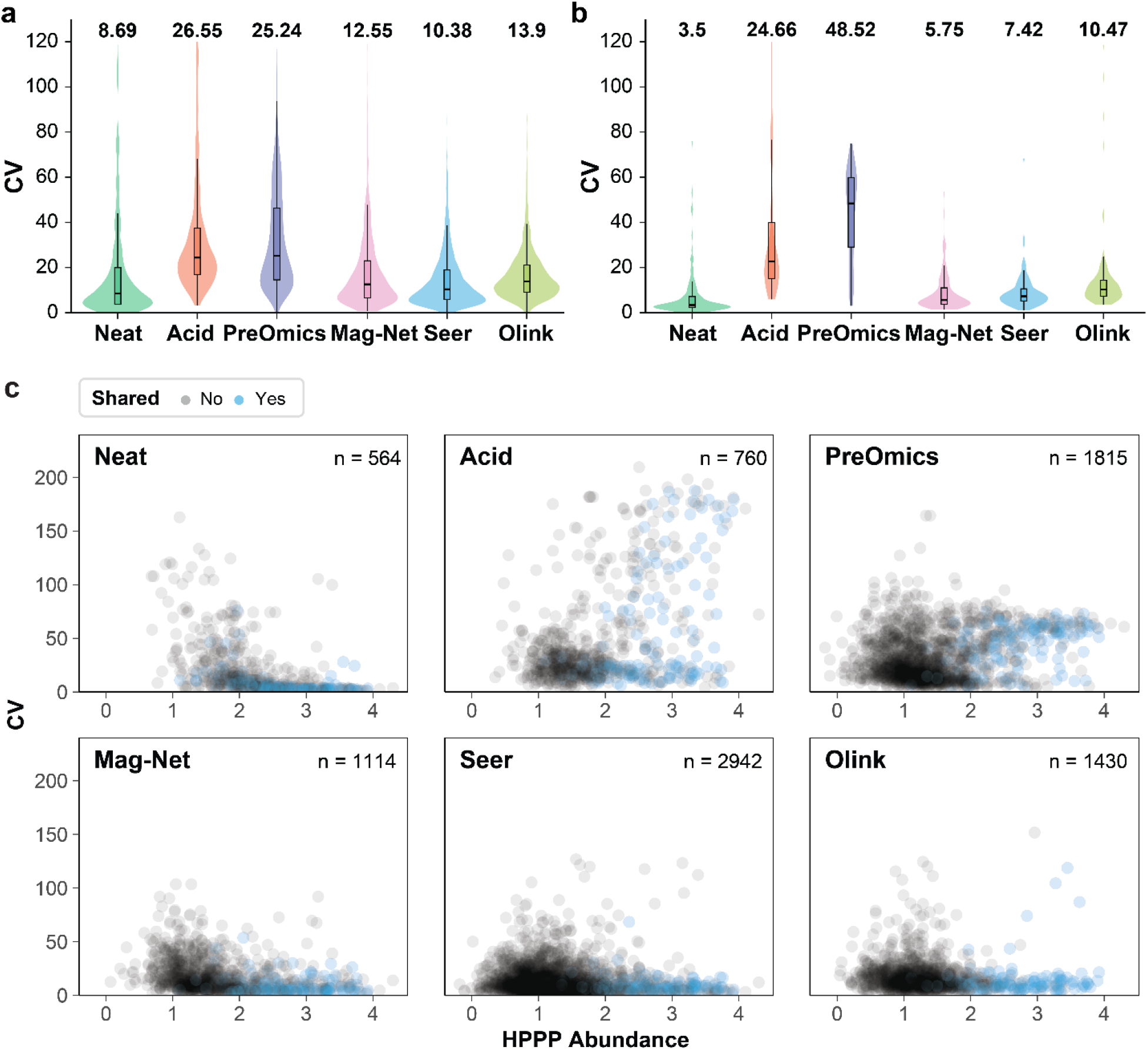
Measurement reproducibility. (a) Coefficient of variation (CV) for protein measurements across five technical replicates. For Olink, protein IDs were only reported for assays that were above LOD. (b) Protein CV for shared protein groups (n = 137) across all six methods. (c) HPPP abundance vs CV for proteins overlapping with HPPP in each method. Shared proteins are shown in blue vs. black dots. Since MS and Olink measurements are not directly comparable, HPPP abundance was used as a proxy for absolute abundance.

To understand the higher CV values for Acid and PreOmics, we examined data from the individual replicates of these two methods and noted the variance in the Acid method was driven primarily by a single sample and for the PreOmics method from two of the samples. In both cases, similar numbers of peptides and proteins were identified; however, the distributions of quant values varied considerably across the datasets (**Supplementary Figure 5**).

### Spike-in standard for linearity assessment

An essential figure of merit for any analytical method is linear response. To determine the linear response of each of the methods, we spiked C-Reactive Protein (CRP) into samples at varying levels, as previously described^20^. First, we measured endogenous CRP levels using ELISA (0.88 μg/mL) and then spiked purified CRP into the sample at 2x, 5x, 10x, and 100x. The spiked samples were split into aliquots and processed in triplicate with each method. Following the same analytical and informatic approaches described above we plotted the measured CRP abundances (**Figure 4)**.

**Figure 4.**
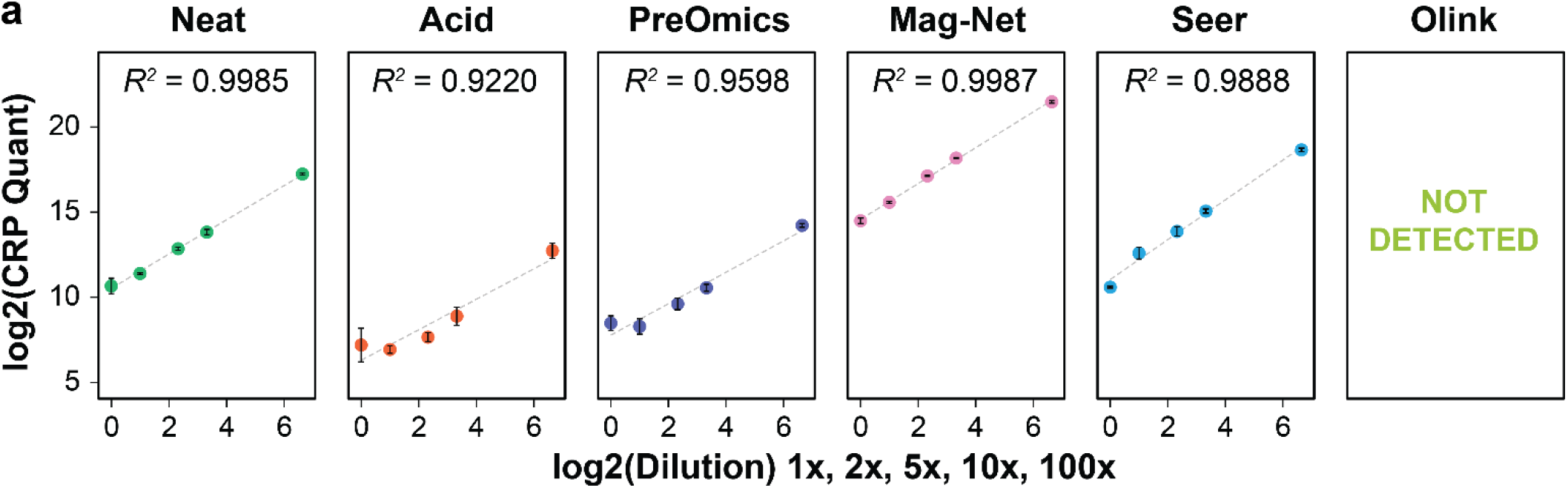
Assessment of measurement linearity with protein spike. Following measurement of endogenous CRP levels, a series of standard additions were made at 2x, 5x, 10x, and 100x the endogenous concentration. Plasma samples were prepared in triplicate at each spike-in level with each method and the abundance of CRP was extracted after analysis using the same protocols and instrument methods as previously described. The measured abundance of CRP was plotted against the spike-in factor.

Notably, given the relevance of CRP in plasma, we did not think to check whether CRP was in the Olink assay and were surprised that it was neither in the assay nor detected. This experiment underscores a disconnect between the affinity-based technologies and discovery proteomics. All the MS-based technologies both detect CRP across samples and perform reasonably well with R^2^ value >0.92. The methods with the highest correlation were Mag-Net at R^2^=0.9987 and Neat at R^2^=0.9985. Both Acid and PreOmics had good linear response at high spike-in values; however, they did not maintain linearity down to endogenous CRP levels. When compared with Neat, the lower quantity in Acid and PreOmics suggest those methods deplete CRP at every addition level.

### Tolerance to elevated lipids

Human plasma can have greatly varying chemical composition, including wide-ranging lipid and metabolite concentrations. Further, recent reports have suggested that high abundance lipid and metabolites could interfere or negatively impact plasma proteomics approaches^36^. To evaluate the robustness of these six methods to potential lipid interference, we spiked triglyceride-rich lipoproteins into plasma at three levels: none, low (100 mg/dL), and high (1,000 mg/dL). The low level is well within normal range for blood triglyceride levels and high is at the extreme end of physiological relevance^37^. The spike-in samples were split and processed once with each method. nLC-MS/MS data was collected with injection replicates (n=3) for each method/spike combination. Olink only had single values for each lipid level due to lack of injection replicates. The number of protein identifications was calculated for each lipid level and method (**Figure 5**). Neat, Acid, Mag-Net, and Seer results were unaffected by the lipid spike-in, while there were subtle differences in the PreOmics method. Additional sample preparation replicates would be required to further explore whether these differences are genuine; however, the small changes may not warrant such effort.

**Figure 5.**
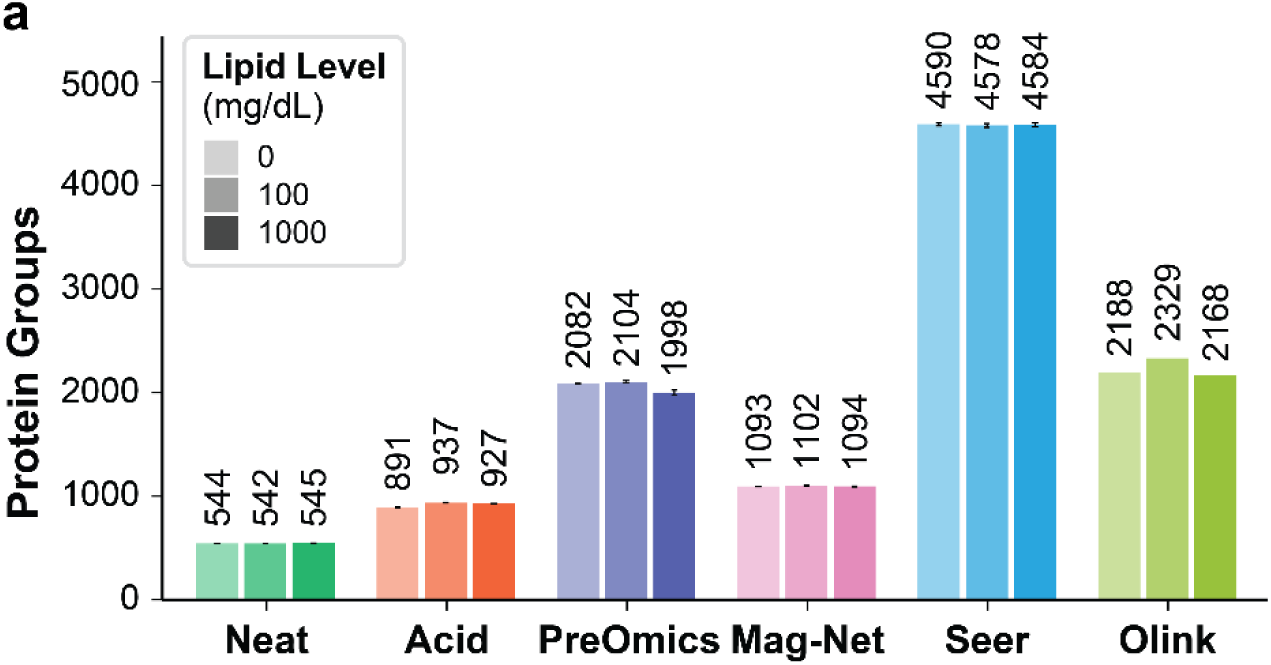
Assessing lipid interference. Triglyceride-rich lipoproteins were spiked into plasma at three levels: control (0 mg/dL), low (100 mg/dL), and high (1000 mg/dL). Plasma samples were prepared once at each spike-in level with each method and measured in injection triplicates with nLC-MS/MS. Olink samples were only measured once. The number of identified protein groups was plotted at each lipid level.

### Global evaluation of protein limit of detection and quantification

While the CRP spike-in experiment provided a window into measurement linearity, it does not provide insight on proteome-wide quantification capability. To globally evaluate the limit of detection and limit of quantitation (LOD and LOQ) for the six plasma proteomics methods in this study we were inspired by Pino et al. and their recent description of a matrix-matched calibration curve (MMCC)^38^. Here we constructed the MMCC by using ten ratios of human plasma diluted in a chicken plasma matrix in triplicate (human:chicken 100:0, 70:30, 50:50, 30:70, 10:90, 5:95, 1:99, 0.5:99.5, 0.1:99.9, 0:100). Each of the six methods was then utilized to process the resulting 30 samples. For information on the MS and Olink data processing, see the Matrix Matched Calibration Curve section in Methods.

**Figure 6** summarizes this data and provides an assessment of global LOD and LOQ for all measured proteins. Because of the filtering performed, some of the proteins for each method will have an infinite LOD/LOQ assigned, due to either variability or few data points, which we labeled as “Detected, not quantified” (**Figure 6a**). Seer had by far the most quantifiable assays for both LOD and LOQ at 4,800, followed by Olink (2,006), PreOmics (1,649), Mag-Net (1,387), Neat (788), and Acid (762). LOD and LOQ results are shown in **Figure 6b** and **6c**. In Figure 6, human plasma dilution LOD/LOQ refers to the percent of human plasma in chicken background where the LOD or LOQ were calculated for each protein. Here we see that Seer produces nearly 2,000 measurments below the 1% human plasma dilution level – which was approximately four-fold higher than the next closest method (PreOmics), with similar trends for the LOQ plot. Interestingly, Olink performs at a level comparable to Neat and Mag-Net (∼300) for the LOD and is the worst performing for LOQ at the 1% human plasma dilution level (25).

We realize that the presence of foreign chicken proten in this experiment presents an atypical challenge for the Olink assay. Even though there is relatively low homology between chicken and human proteins (∼70%), the MS-based methods can eliminate peptides having shared sequences between the two species^39^. Because Olink measures protein abundance through an affinity-based, protein-level assay there is the potential for cross-reactivity between the Olink reagents and chicken protein. To assess the frequency of these events, we evaluated the correlation of the ten human/chicken protein ratios to NPX values and compared that to the same correlations with LFQ values for the MS-based proteomic technologies (**Supplemental Figure 6**). When no Olink suggested LOD cutoff is applied, we see a trimodal distribution with correlations to human proteins, no correlation, and anti-correlation. Filtering by Olink suggested LOD (as we did throughout this work) removes the non-correlated maximum and leaves a bimodal distribution with the majority having positive correlation – supporting that the Olink suggested LOD cutoff should be applied when interpreting Olink data. Finally, we suppose the anti-correlated maximum arises due to cross-reactivity of the Olink antibodies with chicken proteins. Note - we did not use the Olink suggested LOD for our the MMCC calculations as including values below LOD allows noise level measurement and is consistent with the relaxed FDR used in the MS processing.

**Figure 6.**
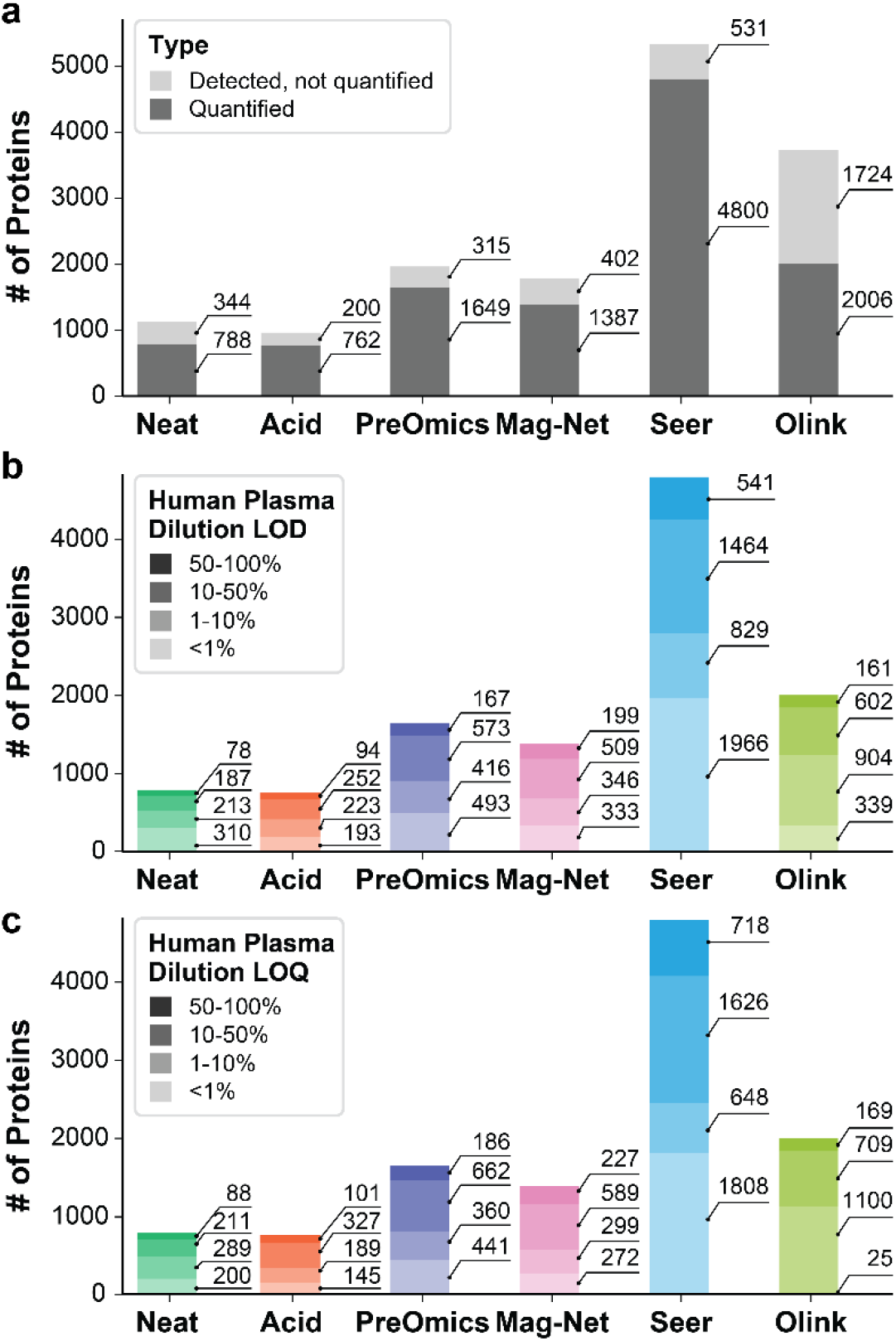
Global evaluation of protein quantification. Human plasma was spiked into a chicken plasma matrix at various levels and the resultant samples processed with each of the methods. (a) Proportion of proteins for each method that could have a LOD or LOQ calculated. (b) For each method, the number of proteins with an LOD calculated at each dilution level with <1% human plasma being the most sensitive. (c) The number of proteins with an LOQ calculated at each dilution level.

### Ability to discover differentially regulated proteins

As a final comparison between the six plasma proteomics methods, we assembled 40 individual plasma samples from two study groups, healthy controls and stage-4 non-small cell lung cancer (NSCLC). Again, samples were analyzed with each method (**Figure 7)**. In a larger biological study, data completeness is an important factor, as too many missing values for a protein across samples limits its utility. **Figure 7a** summarizes the performance of each method with respect to completeness and depth. Seer outperforms the other methods with detection of just over 7,000 proteins, having 6,604 detected in at least half of the samples. PreOmics does well with total identifications (5,157) and 50% completion (4,616); however, this method exhibits the largest gap between 50% and 100% completion with only 1,348 detected in all the samples. Note a small number of the samples had low overall protein identifications (∼2,000) with PreOmics, suggesting potential sample preparation method robustness challenges. Mag-Net ranks next with 3,392 protein groups detected at 50% completeness. These are followed by Olink at 2,199, Acid at 1,728, and Neat at 1,006 proteins quantified at 50% completeness.

**Figure 7.**
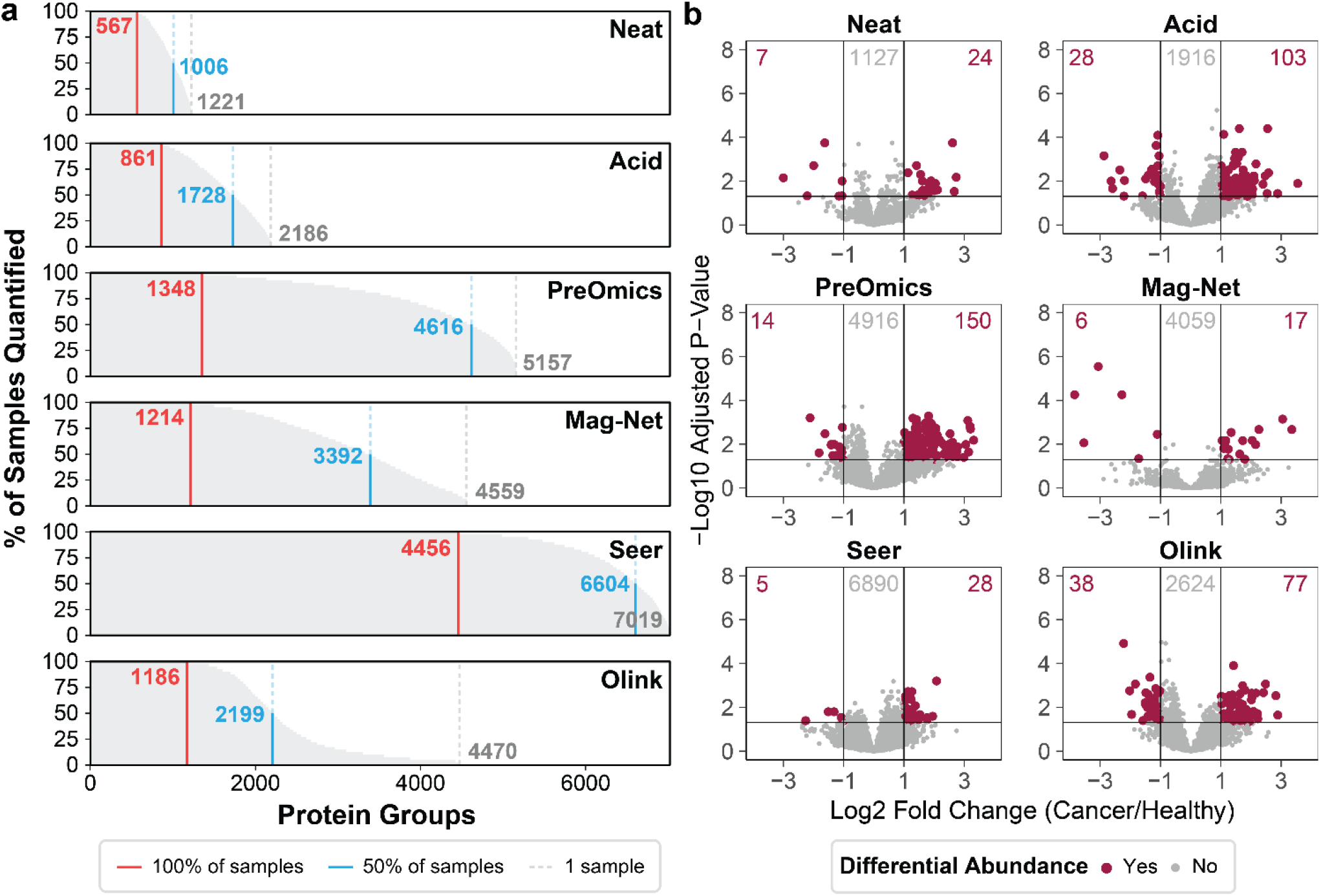
40-sample non-small cell lung cancer (NSCLC) cohort. (a) Summary of identifications and data completeness for each method across the 40 individual plasma samples. Vertical lines indicate the number of proteins identified in 100% of samples, 50% of samples, or at least one sample. (b) Volcano plots of differential abundance between cancer/control groups resulting from each plasma method. Protein abundances were filtered for 80% completeness, log2 transformed, missing values imputed, and t-testing performed. Differential expression was denoted by fold change > 2 and adjusted p-values (BH method) < 0.05.

While depth and completeness are important, evaluating differential abundance of proteins is the primary goal of all these methods. Accordingly, the data for each method was filtered for proteins measured in at least 20% of samples, log2 transformed, and missing values imputed before performing a two-sided t-test between healthy and cancer samples for each protein group. For Olink, protein abundance was additionally normalized by sample to the median NPX value, as there was no control for overall protein content (**Supplemental Figure 7**) whereas for the MS-based methods peptide loading mass was the same for every sample. Details on data analysis can be found in the Differential Expression Calculations section in Methods. Differentially expressed protein groups for each plasma method are presented in **Figure 7b**. Using these basic statistical tests, all the methods find significant differences between the healthy and cancer groups. However, PreOmics and Acid generate more differentially abundant proteins than any of the other methods. Further, all the methods, but especially those two, have strong bias to detect up-regulated proteins. We suppose this could be explained by the need to control for co-variates (i.e., age, sex, severity, etc.) and/or more sophisticated statistical testing. One may also expect that cancer would increase the presence of proteins that aren’t normally present in plasma (i.e., cell leakage, etc.). Finally, it seems likely that measurement reproducibility across the methods – especially when data is imputed – would play into the number of significant findings.

## Conclusion

Here we present a direct comparison of six methods for plasma proteomics using multiple analytical figures of merit. Methods were assessed for achievable proteomic depth, reproducibility, linearity of measurement, robustness to lipid interference, global limit of detection and quantification, and use in a differential expression study. **Table 1** both summarizes the results and provides other relevant method comparisons.

**Table 1.** Summary of plasma proteomics technologies. The values presented in this table represent this study and the implementation used herein. *Automation is available for all methods, but only implemented for Seer and Olink here. ^†^The CRP protein was not included in the Olink assays so we could not assess linearity for that protein. ^‡^The use of chicken plasma for the matrix matched calibration curve presents a unique challenge in estimated LOD/LOQ for Olink.

| Method | Starting Plasma (μL) | Samples /day | Cost | Automation* | Prep Time (overall / person hours) | Approximate Depth (Proteins) | Median Protein CV (n = 5) | CRP Linearity† | MMCC LOD/LOQ‡ |
| --- | --- | --- | --- | --- | --- | --- | --- | --- | --- |
| Neat | 5 | 24 | \$ | No | 24 / 3 | 600 | 8.7 | ★★★★★ | ★★★ |
| Acid | 20 | 24 | \$ | No | 24 / 3 | 800 | 26.5 | ★★★ | ★★★ |
| PreOmics | 50 | 24 | \$\$ | No | 6 / 4 | 2,200 | 25.2 | ★★★ | ★★★ |
| Mag-Net | 100 | 24 | \$ | No | 8 / 4 | 1,200 | 12.5 | ★★★★★ | ★★★ |
| Seer | 240 | 12 | \$\$\$ | Yes | 9 / 2 | 4,500 | 10.4 | ★★★★ | ★★★★★ |
| Olink | 10 | 172 | \$\$\$ | Yes | 1 / 1 | 2,600 | 13.9 | NA | ★★ |

Considering **Table 1**, the recommended plasma starting volume ranges from <5 μL for Neat up to 240 μL per sample for Seer. We note that high sample volumes could prohibit certain applications, i.e., small mammal research; however, the affinity technology Olink, which only requires ∼ 10 μL, is designed specifically for human proteins. Samples per day is a key consideration for many plasma studies. The MS-based methods all, except for Seer, could achieve a throughput of ∼ 24 samples per day. In this study throughput was limited by the time required for MS data acquisition, which we optimized for performance rather than speed. This will continue to improve as technology matures but at present it is the primary bottleneck. The Olink approach scales much higher owing to its reliance on next-generation sequencing technology.

Cost is another key consideration and here and several MS methods including Neat, Acid, and Mag-Net had minimal reagent costs. PreOmics (pre-released ENRICHplus) and Seer Proteograph XT are both commercial kits and those technologies incur additional cost. The MS methods also incur the cost of operating an LC-MS system, which for the majority of these methods comprises the bulk of the overall per-sample expense. Currently Olink has a flat cost per sample, which is higher than most of the MS-based methods and comparable to Seer. In terms of sample preparation, both overall and hands-on time varied across the methods. Since Olink is a service, the main labor spend was on aliquoting the samples for shipment. Seer also had minimal sample preparation given that the procedure is performed on an automated system integral to the protocol. Neat and Acid were straightforward with a limited number of steps. The kit-based format of PreOmics was easy to follow, but the number of washing steps for both PreOmics and Mag-Net increased hands-on time significantly.

The final section of **Table 1** covers the technical performance of the various approaches, which we discuss by method. Neat offered the least proteomic depth (600); however, it had the best precision and excellent quantitative linearity with moderate LOD/LOQ. The Acid depletion method slightly improved depth but at the cost of considerably reduced precision, reduced linearity, and had moderate LOD/LOQ. PreOmics greatly increased proteomic depth of Neat and Acid but also suffered from high variability (25.2%), and had moderate performance for linearity and LOD/LOQ. The next method, Mag-Net, had a two-fold boost in protein depth over Neat, excellent linearity and precision with average LOD/LOQ. Seer offered the best proteomic depth (∼4,500), excellent quantitative precision and linearity, and the best LOD/LOQ observed in the study. Olink had the second-best proteomic depth, slightly better than PreOmics with 2,600 proteins, good quantitative precision, but was worst in terms of LOD/LOQ.

Overall, this data demonstrates there are many technical details to consider when selecting a technology to profile plasma proteins. First, we conclude that among the MS-based technologies, the method by which one prepares the sample for MS analysis strongly influences the performance – especially the depth. When compared to Olink, the best performing MS technology, Seer, offers nearly double the depth with stronger technical figures of merit in every category. But beyond these benefits, the MS-based approaches offer a true discovery platform wherein the Olink strategy is best described as a targeted proteomic technology. MS-based data has the further benefit, especially with DIA analysis, for future mining of PTMs, variants, and the presence of proteins from other organisms, i.e., bacteria or even chicken. On the other hand, Olink has high sample throughput owing to its use of next-generation sequencing technologies, with published studies upward of 50,000 sample cohorts. Another interesting result here was that the overlap between the MS and Olink identifications was minimal, providing complementarity as previously reported.

## Acknowledgements

We thank the NIH grant P41 GM108538 and R35 GM118110 for financial support of this work. We also thank Seer, PreOmics, Olink, and the Steen laboratory. Specifically, Seer provided materials (Xiaoyan Zhao: kits, reagents, and purchased plasma), technical advice (Asim Siddiqui) and analysis support (Lee Cantrell, Ting Huang, Jian Wang: matrix-matched calibration curve and differential expression analysis), and sponsored the Olink data collection. PreOmics provided access to the pre-release EnrichPLUS plasma kit and on-site technical support (Measho Abreha). We are also grateful for technical support from Olink and helpful discussions with Dylan Tabang of the Hanno Steen laboratory (Acid). Note that JJC is on the Scientific Advisory Board of Seer. He is a consultant for Thermo Fisher Scientific and a founder of CeleramAb Inc.

## Methods

### Experimental setup

Experiment 1 consisted of five identical plasma samples prepared in tandem with each method. Experiment 2 consisted of five spike-in levels of C-Reactive Protein (CRP) prepared in triplicate with each method. Experiment 3 consisted of three spike-in levels of triglyceride-rich lipoproteins prepared and then analyzed in triplicate injections with LC-MS/MS (or once with Olink). Experiment 4 consisted of a matrix-matched calibration curve, described below. Experiment 5 was a pilot biological study constructed of 20 plasma samples from stage 4 non-small cell lung cancer (NSCLC) patients and 20 age- and sex-matched healthy control samples prepared with each method. Plasma for experiment 5 was also obtained from BioIVT. Samples were randomized before LC-MS data collection. The Seer Proteograph XT kit divides each sample into an NPA and NPB fraction, so to analyze of a sample via Proteograph XT used 2 hours of LC-MS/MS time vs 1 hour for all other methods.

### Plasma Aliquoting

Frozen K2EDTA plasma was purchased from BioIVT (cat. HUMANPLK2-0101355). Plasma was thawed once, centrifuged at 3000*g* for 5 min, prepared with appropriate spike-ins and dilutions, aliquoted, and refrozen in Eppendorf Lo-Bind tubes. For the C-Reactive Protein (CRP) Spike-In, CRP was obtained (Sigma Aldrich, cat. AG723-M) and prepared in standard dilutions from 1 mg/mL to 0.01 mg/mL. Volume of standard or milli-Q water were added to equal volumes of plasma for each spike in to ensure the same relative concentration of proteins in plasma. For the lipid spike in, a triglyceride-rich lipoprotein standard was obtained (Sun Diagnostics, cat. INT-01T). Either milli-Q water or lipid standard were added at equal volumes to maintain a constant dilution of the plasma. For the matrix matched calibration curve, the standard human plasma was diluted in K2EDTA chicken plasma background at 10 different ratios to create a calibration curve (human:chicken 100:0, 70:30, 50:50, 30:70, 10:90, 5:95, 1:99, 0.5:99.5, 0.1:99.9, 0:100). Before mixing volumetrically, the total protein content of both types of plasma was measured by protein BCA assay, human plasma was 71 mg/mL and chicken plasma was 51 mg/mL. The human plasma was diluted with milli-Q water to match protein concentration of chicken before mixing. The ratios were each prepared individually and not serially to prevent cascading effects.

### Neat Plasma Preparation

Neat plasma preparation was based on the method from Shishkova and Coon, 2021^33^. In Briefly, 5 μL of plasma was mixed with 50 μL Lysis Buffer A (4.8 M Guanidine HCl (Sigma-Aldrich, G3272), 10 mM TCEP (Sigma-Aldrich, C4706-2G), 40 mM 2-chloroacetamide (Sigma-Aldrich, ≥98%, C0267-100G), 100 mM Tris pH 8 (Invitrogen, 1 M Tris pH 8.0, 0.2 μm filtered, AM9856)) and heated at 100 °C for 10 min. Proteins were precipitated with addition of 450 μL methanol (Optima LC/MS grade, Fisher Scientific), vortexed, pelleted, and air dried after decanting supernatant. Pellets were resuspended in 100 μL Lysis Buffer B (5.33 M Urea (Sigma-Aldrich, U5378), 5mM TCEP, 20 mM 2-chloroacetamide, 100 mM Tris pH 8) until fully resolubilized. Samples were diluted to 1.2 M urea with additional 100 mM Tris pH 8. 5 μg Lysyl Endopeptidase (LysC, 100369-826, VWR) and 7.5 μg trypsin (Promega, V5113) per sample were added for overnight digestion at RT. Digested samples were acidified to ∼1% trifluoroacetic acid by addition of 10% trifluoroacetic acid (Sigma-Aldrich, HPLC grade, >99.9%).

Samples were desalted with 10 mg Strata-X 33 μm polymeric reversed phase SPE cartridge (Phenomenex, 8B-S100-AAK). Primed with 1 mL acetonitrile (Optima LC-MS grade, Fisher Scientific) and 1 mL 0.1% trifluoroacetic acid, samples loaded, washed with 1 mL 0.1% trifluoroacetic acid, then eluted with 300 μL 80% acetonitrile/0.1% trifluoroacetic acid and dried down in a SpeedVac (Thermo Scientific) and stored at -80 °C until reconstitution.

### Perchloric Acid Depletion

Perchloric acid protein depletion was performed based on the protocol from Viode et al. 2023^22,23^. Briefly, 20 μL plasma was added to a 1.5 mL microfuge tube with 480 μL milli-Q water. Added 25 μL 70% perchloric acid (Sigma-Aldrich, 244252-100ML) and vortexed for 3 min. Cooled samples at -20 °C for 15 min. Centrifuged samples for 60 min at 3200*g*, 4 °C to pellet precipitated proteins. Samples were acidified by taking supernatant (400 μL) and adding 40 μL of 1% trifluoroacetic acid (Sigma-Aldrich, HPLC grade, >99.9%). An Oasis μSPE HLB plate (Waters, 186001828BA) was prepared using a vacuum manifold. Plate was conditioned using 300 μL of methanol (Optima LC/MS grade, Fisher Scientific) and 2x 500 μL 0.1% trifluoroacetic acid. Samples were loaded and washed 3x with 500 μL 0.1% trifluoroacetic acid. Proteins were eluted from the plate with 100 μL 90% acetonitrile/0.1% trifluoroacetic acid into a lo-bind 0.5 mL plate (Eppendorf, 951032107) and dried in a SpeedVac (Thermo Scientific). Samples were reconstituted with 42.5 μL 50mM ammonium bicarbonate (Sigma-Aldrich, A6141) and digested with 1 μg trypsin (Promega, V5113) for 18 h at 37 °C. Digested peptides were acidified with 5 μL 10% formic acid (Fisher Scientific, LC-MS grade). Samples were desalted with 10 mg Strata-X 33 μm polymeric reversed phase SPE 96-well plate (Phenomenex, 8E-S100-AGB). Primed with 1 mL acetonitrile (Optima LC-MS grade, Fisher Scientific) and 1 mL 0.1% trifluoroacetic acid, samples loaded, washed with 1 mL 0.1% trifluoroacetic acid, then eluted with 300 μL 80% acetonitrile/0.1% trifluoroacetic acid and dried down in a SpeedVac (Thermo Scientific) and stored at -80 °C until reconstitution.

### PreOmics ENRICHplus

50 μL of plasma was prepared using the pre-release ENRICHplus kit according to manufacturer’s protocol (PreOmics, Planegg/Martinsried, Germany). All steps were performed by hand in a 96-well plate format using lo-bind 0.5 mL plates (Eppendorf, 951032107) with sealing mats (Eppendorf, 0030127978). First, 200 μL ENplus-WASH was added to each well and mixed with 25 μL ENplus-BEADS (beads were thoroughly resuspended before adding to wells), then plate was sealed and mixed at 1200 rpm for 1 min using a Multitherm Heatshake (Benchmark Scientific). Beads were pulled down with a magnetic rack (Magnum EX, Alpaqua) and supernatant discarded. Washing was repeated 2x more for a total of 3 washes. 50 μL plasma and 50 μL ENplus-BIND buffer were added to each well and mixed for 30 min at 1200 rpm and 30 °C. The plate was placed on a magnetic rack and the unbound components discarded. To wash, 100 μL ENplus-BIND was added to each well and mixed at 1200 rpm for 1 min before placing the plate on the magnetic rack and discarding the supernatant. Washing was repeated 2x more for a total of 3 washes. 50 μL LYSE-BCT was added to each well and mixed for 10 min at 1200 rpm and 60 °C before cooling to RT. 50 μL DIGEST was added to each well and mixed at 1200 rpm and 37 °C for 1 h. 100 μL STOP was added to each well and mixed to end digestion. To purify samples, they were transferred to elution cartridges, then washed with 200 μL WASH 1 and 200 μL WASH 2 before eluting into the collection plate with 200 μL ELUTE. The collection plate was then centrifuged, and 180 μL of the elutant was transferred to a new plate to dry down in a SpeedVac (Thermo Scientific) and stored at -80 °C until reconstitution.

### Mag-Net^21^

The Mag-Net method was performed generally according to the method detailed on the ReSyn website (Version 5, Dec. 2023). All steps were performed by hand in a 96-well plate format using lo-bind 0.5 mL plates (Eppendorf, 951032107). Briefly, for each sample 25 μL MagReSyn SAX beads (ReSyn, MR-SAX005) were equilibrated 2x in 200 μL BTP Equilibration/Wash Buffer (50 mM Bis Tris propane pH 6.4 (Sigma-Aldrich, B6755), 150 mM sodium chloride (Fisher Scientific, S271-1)). The beads were held with a magnetic rack (Magnum EX, Alpaqua) and supernatant decanted, then 100 μL bind buffer (100 mM Bis Tris propane pH 6.4, 150 mM sodium chloride) and 100 μL plasma were added to the beads and mixed at 900 rpm for 30 min on a Multitherm Heatshake (Benchmark Scientific). The magnetic beads were held with a magnetic rack and supernatant decanted, then washing was repeated 3x. To wash, 400 μL BTP Equilibration/Wash Buffer was added to each sample, mixed for 5 min at 900 rpm, then beads held with a magnetic rack while supernatant was discarded. After washing, beads were resuspended in 100 μL lysis and reduction mix (50 mM Tris pH 8 (Invitrogen, AM9856), 1% (w/v) SDS (Sigma-Aldrich, L4509), 10 mM TCEP (Sigma-Aldrich, C4706-2G), 40 mM 2-chloroacetamide (Sigma-Aldrich, ≥98%, C0267-100G)) and incubated for 60 min at 37 °C and 900 rpm. To induce on-bead precipitation, 240 μL acetonitrile was added to each well, pipetted up and down, and incubated at RT for 10 min. Plate was moved to the magnetic rack and supernatant removed. With the plate on the rack, 95% acetonitrile was added, incubated for 30 s, and removed. Washing was repeated 2 more times for a total of 3 organic washes. For digestion, beads were resuspended in 200 μL digestion mixture (50 mM ammonium bicarbonate (Sigma-Aldrich, A6141), 7.5 μg trypsin (Promega, V5113), 2 μg Lysyl Endopeptidase (LysC, 100369-826, VWR)) and incubated at 47 °C for 2 h at 900 rpm mixing. Digestion was quenched with 12.5 μL 10% trifluoroacetic acid and the peptide containing supernatant removed for further processing. Samples were desalted with 10 mg Strata-X 33 μm polymeric reversed phase SPE 96-well plate (Phenomenex, 8E-S100-AGB). Primed with 1 mL acetonitrile (Optima LC-MS grade, Fisher Scientific) and 1 mL 0.1% trifluoroacetic acid, samples loaded, washed with 1 mL 0.1% trifluoroacetic acid, then eluted with 300 μL 80% acetonitrile/0.1% trifluoroacetic acid and dried down in a SpeedVac (Thermo Scientific) and stored at -80 °C until reconstitution.

### Seer Proteograph XT

240 μL of plasma was prepared using the Proteograph XT kit according to manufacturer’s protocol (Seer Inc., Redwood City, USA). With the SP100 automated liquid-handling robot, samples and controls were distributed into a 96-well plate. Each plasma sample was incubated with 2 nanoparticle mixtures to create 2 fractions per sample (NPA and NPB). After nanoparticle incubation and corona formation, nanoparticles were held using a magnetic rack unbound components removed with a series of washes. Bound proteins were processed through reduction, alkylation, and digestion with trypsin/LysC before elution into a collection plate using a positive pressure system (MPE). Peptides were quantified using the Pierce Quantitative Fluorometric Peptide Assay (Thermo Scientific, 23290) before being dried down in a SpeedVac (Thermo Scientific) and stored at -80 °C until reconstitution.

### LC-MS Data Collection

Dried peptides from each method were resuspended in 0.2% formic acid (Fisher Scientific, LC-MS grade) between 0.05 and 1 μg/μL and quantified either with the Quantitative Fluorometric Peptide Assay (Thermo Scientific, 23290) for Seer Proteograph XT or Pierce Quantitative Colorimetric Peptide Assay (Thermo Scientific, 23275) for all others. For each method 300 ng peptides were loaded for analysis (between 0.2-5 μL loading volume).

Peptides were separated using a Vanquish Neo nanoLC system (Thermo Scientific). For separation, a 75 μm inner diameter nanoflow capillary column was packed to 45 cm with 1.7 μm, 130 Å pore size BEH C18 particles (Waters) in-house.^34^ Briefly, a slurry of 11 mg C18 particles suspended 110 μL chloroform was loaded into a custom-built packing setup with a high pressure pneumatic pump (Haskel) and packed into the column while slowly ramping up to 25000 psi, holding 1 h, and then allowed to depressurize naturally. The column was coupled to the mass spectrometer with a NanoSpray Flex source set to 2100 V (Thermo Scientific) and heated to 50 °C with a Column Oven PRSO-V2 (Sonation Lab Solutions). Mobile phase A was 0.2% formic acid in water, and mobile phase B was 80% acetonitrile/20% 0.2% formic acid in water. The gradient elution was carried out at 0.300 μL/min with a 30-minute active gradient. Mobile phase was ramped from 0%B to 10%B from 0 to 2 min. The active gradient was set to 10% to 54%B with curve type 6 beginning at 2 min, adjusted to evenly distribute peptide signal across the gradient. The column was washed for 7 min at 100%B at the end of the gradient, followed by column equilibration at 0%B.

Mass Spectrometry analysis was performed using positive mode Data-Independent Acquisition (DIA) on the Orbitrap Astral MS. MS^1^ were acquired in Orbitrap mode with 240k resolution every 0.6 s over 380-980 *m/z*. The MS^1^ normalized AGC target was set to 250% (2.5e6 charges) with a maximum injection time of 50 ms and an RF Lens setting of 40%. Astral DIA MS^2^ were acquired with a precursor isolation range of 380-980 *m/z* divided into 4 *m/z* DIA windows with window placement optimization on, resulting in 150 MS^2^ scans per cycle. Normalized AGC target was set to 100% (1e4 charges) with a 5 ms max injection time. HCD collision energy was set to 25% with a default charge state of +2. If all MS^2^ scans hit max injection time the MS^2^ max cycle time would be 0.75 s. Instrument QCs were 100ng neat plasma peptides to monitor performance.

### LC-MS Data Analysis

LC-MS data for experiments 1-3 were processed using separate dDIA searches for each method/experiment combination on Spectronaut 19 with default settings. Seer NPA and NPB were searched together. Data for experiments 1-3 and 5 were analyzed with RStudio. Spectronaut reports were filtered for 0.01 PG.Qvalue and 0.01 EG.Qvalue while PG.Quantity was used for quantitative comparisons. Seer NPA and NPB were combined into a collective unit via maximum representation (iterating through each protein group and choosing which NP had greater data completeness).

### Olink Explore HT

The relative abundance of 5415 plasma proteins was determined using the Olink Explore HT Platform at Life & Brain GmbH Olink certified laboratory in Bonn, Germany (Olink Proteomics, Uppsala, Sweden). Protein abundance in 10 μL of plasma was quantified. Briefly, relative protein abundance in plasma was quantified based on the binding of pairs of oligonucleotide-labeled antibodies to each protein. The barcodes are then read with a NovaSeq 6000 sequencing system (Illumina, San Diego, CA, USA). Olink proteins were reported on a relative and log2 scale as Normalized Protein eXpression (NPX) values. In Experiments 1-3 and 5, measurements were filtered for only those that passed Assay QC and Sample QC and were above LOD as calculated by the OlinkAnalyze R package^40^. Values below LOD were reported as missing values. For the MMCC in Experiment 4, all values were included as a measure of noise.

### Matrix Matched Calibration Curve

Total protein content was measured for human and chicken plasma, human was diluted in water to match the concentration of chicken. 10 dilutions were made volumetrically combining the 2 types of plasma (human:chicken 100:0, 70:30, 50:50, 30:70, 10:90, 5:95, 1:99, 0.5:99.5, 0.1:99.9, 0:100). These dilutions were prepared in triplicate for each method.

MS samples were run in order of increasing concentration of human plasma. Raw files were searched separately for each method using DIANN 1.8.1 in library-free mode. The searches used the mouse reference proteome (both Swiss-Prot and TrEMBL) downloaded from UniProt on 08-Oct-2024 and the human reference proteome (only Swiss-Prot) downloaded from UniProt on 06-Aug-2024. DIANN was set to filter at 50% FDR for measurement of effective noise. Each report was imported into R and filtered for 0.01 Lib.Q.Value. Next, only uniquely human peptides were selected for protein grouping and then normalized using maxLFQ. Protein groups were filtered for >10% completeness and LOD and LOQ were calculated for each protein group in each method according to the statistical analysis described in Pino et al. 2020^38^. Code for generation the LOD and LOQ was found on GitHub (https://github.com/sjust-seerbio/matrix-matched-calcurves). Infinite LOD and LOQ values were managed according to **Table 2**.

**Table 2.** Procedure for infinite values generated during LOD/LOQ calculation for MS-based methods.

|  |
| --- |
| LOD |
| is.infinite(stddev_noise) & slope_linear == 0 & is.infinite(LOD) ~ Inf |
| is.infinite(stddev_noise) & slope_linear > 0 & is.infinite(LOD) ~ 0 |
| is.infinite(LOD) == F ~ LOD |
| LOQ |
| is.infinite(LOD) ~ Inf |
| is.infinite(LOD) == F & is.infinite(LOQ) & slope_linear > 0 ~ LOD |
| is.infinite(LOD) == F & is.infinite(LOQ) == F ~ LOQ |
| LOD == 0 & is.infinite(LOQ) & slope_linear > 0 ~ 0 |
| LOD == 0 & is.infinite(LOQ) & slope_linear == 0 ~ Inf |

For the Olink MMCC, LOD and LOQ were calculated both in the forward and reverse dilution direction to evaluate response to any interfering chicken proteins. Infinite values for either chicken or human were assessed using the arguments in **Table 3** and proteins specific for each were stored. We additionally calculated three metrics for assay measurability using the Olink negative controls (*NPX* > *Mean*_*neg*_ + (3 ∗ σ_*neg*_), *NPX* > *Mean*_*neg*_ + (3 ∗ *MAD*_*neg*_), *NPX* > *Olink LOD*). Chicken or human responding assays were considered that had at least one replicate above Olink suggested LOD (Measurement Reproducibility > 0) (**Supplemental Figure 8**). Once the assays were chosen and sorted into either human measurable or chicken measurable by the stored protein lists, they were categorized into different groups of ‘Chicken Measured’ ‘Human Measured’ ‘Interfering Signal’ Non ‘Definite Grouping’ or ‘Unmeasurable Signal’ according to the arguments in **Table 4**. The number of assays falling into each grouping category is shown in **Table 5**.

**Table 3.** Procedure for infinite values generated during LOD/LOQ calculation for Olink.

|  |
| --- |
| LOD |
| is.infinite(stddev_noise) & slope_linear <= 0.000 & is.infinite(LOD) ~ Inf |
| is.infinite(stddev_noise) & slope_linear > 0 & is.infinite(LOD) & cor >= 0.6 ~ 0 |
| is.infinite(stddev_noise) & slope_linear > 0 & is.infinite(LOD) & cor < 0.6 ~ Inf |
| is.infinite(LOD) == F ~ LOD |
| LOQ |
| is.infinite(LOD) ~ Inf |
| is.infinite(LOD) == F & is.infinite(LOQ) & slope_linear > 0.001 ~ LOD |
| is.infinite(LOD) == F & is.infinite(LOQ) == F ~ LOQ |
| LOD == 0 & is.infinite(LOQ) & slope_linear > 0.000 ~ 0 |
| LOD == 0 & is.infinite(LOQ) & slope_linear <= 0 ~ Inf |

**Table 4.** Grouping for Olink assays after combining LOD/LOQ calculations from forward and reverse dilution directions.

|  |
| --- |
| 'CHICK Response' & proportion_measured > 0 ~ 'Chicken Measured' |
| 'CHICK Response' & proportion_measured == 0 ~ 'Interfering Signal' |
| 'HUMAN Response' & proportion_measured > 0 ~ 'Human Measured' |
| 'HUMAN Response' & proportion_measured == 0 ~ 'Interfering Signal' |
| 'Not Measurable' & proportion_measured > 0 & proportion_measured < 1 ~ 'Non Definite Grouping' |
| 'Not Measurable' & proportion_measured == 0 ~ 'Unmeasurable Signal' |
| 'Not Measurable' & proportion_measured == 1 ~ 'Interfering Signal' |

**Table 5.**
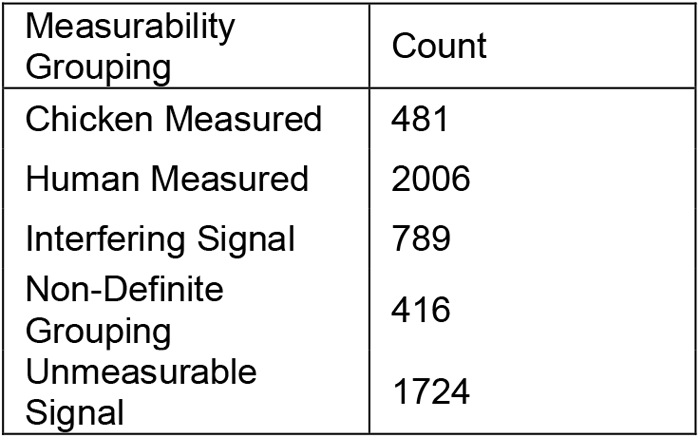
Final grouping of Olink assays from the MMCC experiment.

| Measurability Grouping | Count |
| --- | --- |
| Chicken Measured | 481 |
| Human Measured | 2006 |
| Interfering Signal | 789 |
| Non-Definite Grouping | 416 |
| Unmeasurable Signal | 1724 |

### Differential Expression Calculations

For differential expression analysis between the 20 healthy and 20 cancer samples for each method, first protein quantities were log2 transformed to normalize the data. Proteins with missing measurements from >= 80% of samples were excluded. Then data was imputed using a method similar to Perseus^41^. By column, missing values were imputed from a normal distribution down shifted 1.8 standard deviations from main distribution with a standard deviation width of 0.3. Olink data was median normalized (column-wise). A two-sided t-test was performed between the healthy and cancer groups for each protein group. Differentially expressed protein groups were defined as those with an FC > |2| and *p* < 0.05 using Benjamini–Hochberg multiple testing correction.

