## Supplemental Figures for "A Technical Evaluation of Plasma Proteomics Technologies"

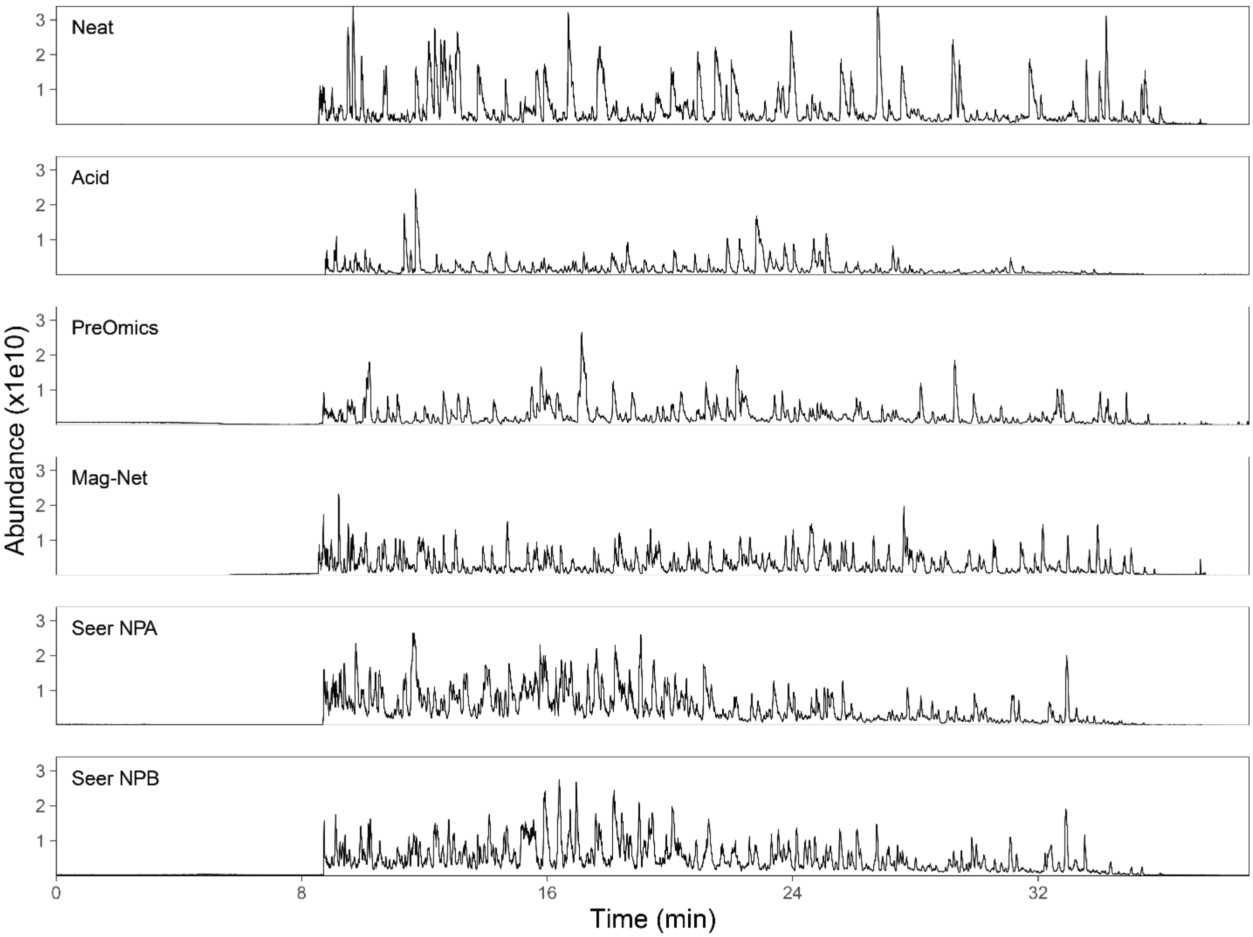


**Supplemental Figure 1 | Representative chromatograms.** Total ion current chromatograms from 300 ng peptide loading mass for each method over a 39-minute LC gradient.

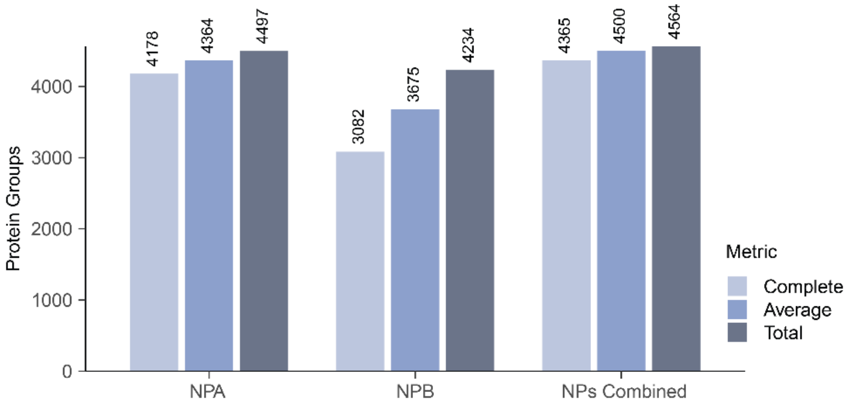


**Supplemental Figure 2 | Comparison of protein identifications for Seer Proteograph XT nanoparticles.** Protein identifications for technical replicates (n = 5) of individual nanoparticle fractions and combined fractions. “Complete” is the number of protein groups detected in all five replicates, “Average” is the average number of protein groups detected per replicate, and “Total” is all unique protein groups detected in at least one replicate.

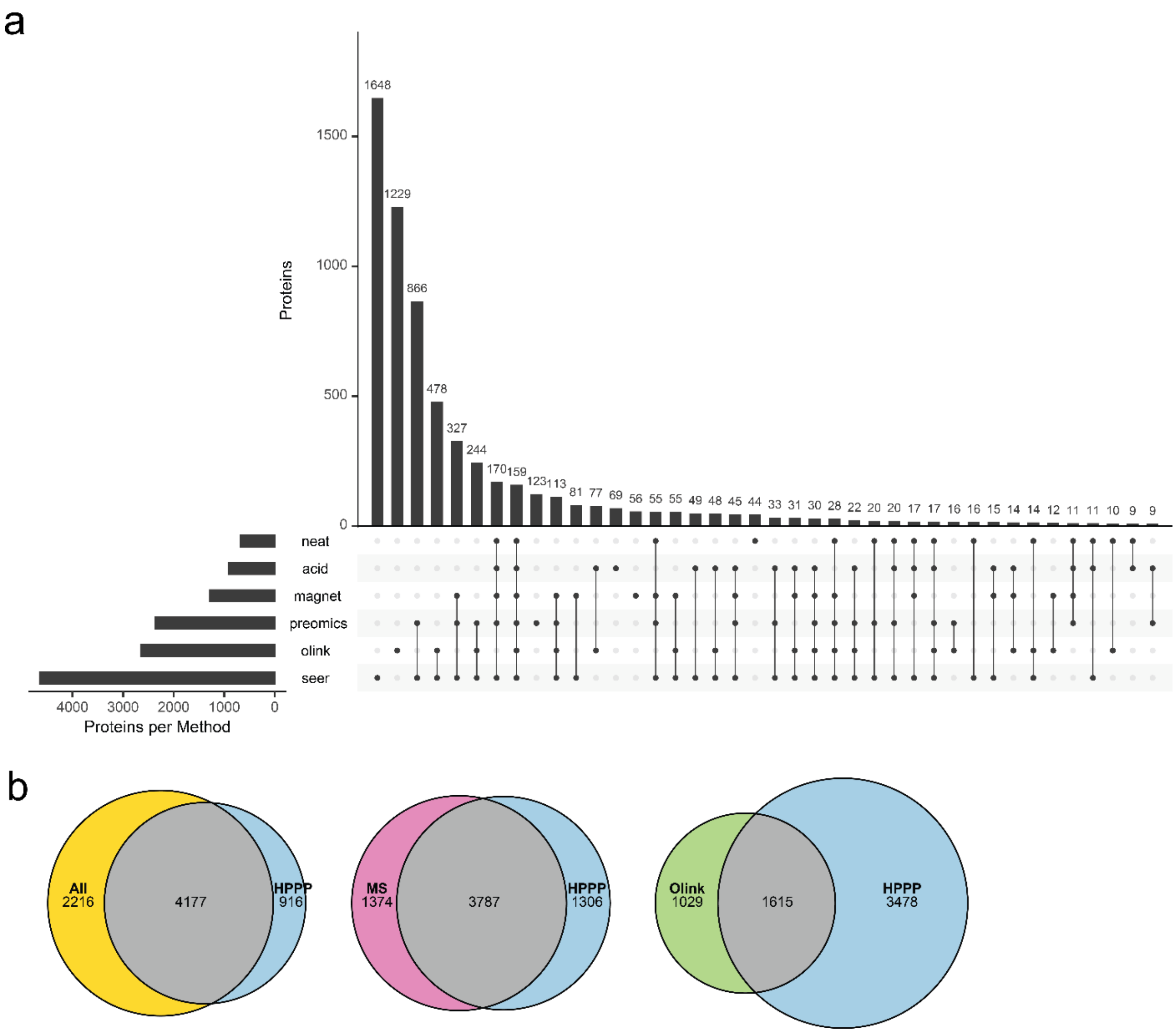


**Supplemental Figure 3 | Identification overlap among plasma methods and Human Plasma Proteome database.** (a) UpSet plot representing gene-level protein identification overlap among all methods in this study. (b) Gene-level overlap between groups of proteins from this study and the 2021 Human Plasma Proteome database. The “All” group represents proteins identified in at least one method in this study, “MS” group contains the MS-based methods.

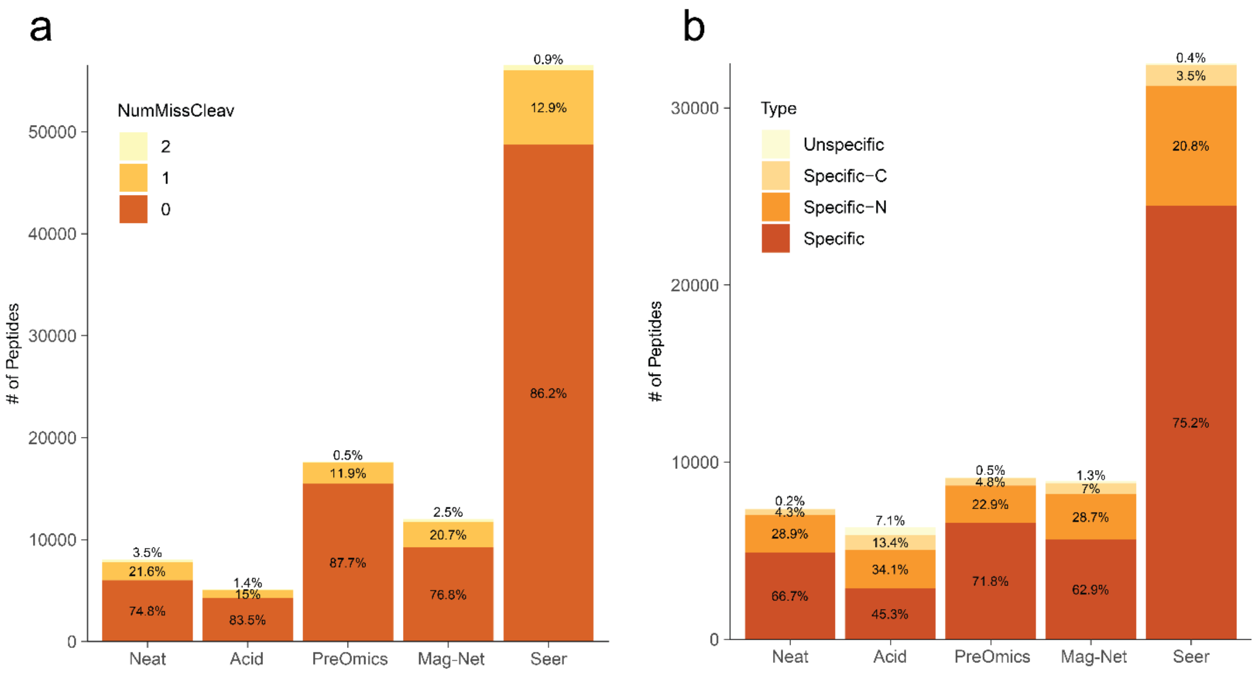


**Supplemental Figure 4 | Missed and unspecific cleavages.** (a) Percent of peptides with missed tryptic cleavages calculated for each MS plasma proteomics method. (b) Percentage of unspecific or specific tryptic cleavages for each method.

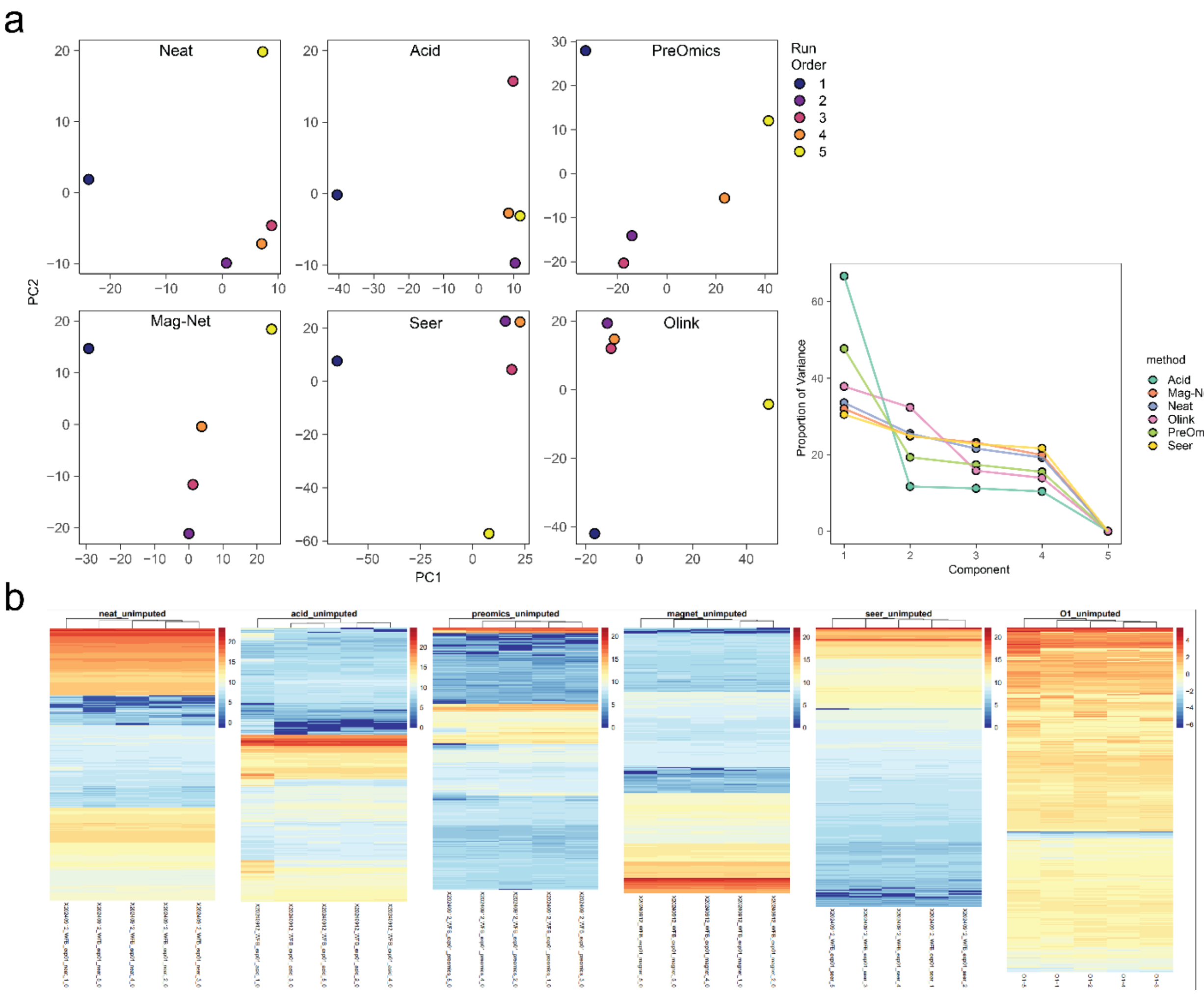


**Supplemental Figure 5 | Variation among protein quantification in experiment 1.** (a) Principal Component Analysis of samples in experiment 1 representing variance in protein quantification. Samples are colored by run order. (b) Heatmaps of protein quantification across the technical replicates for each method.

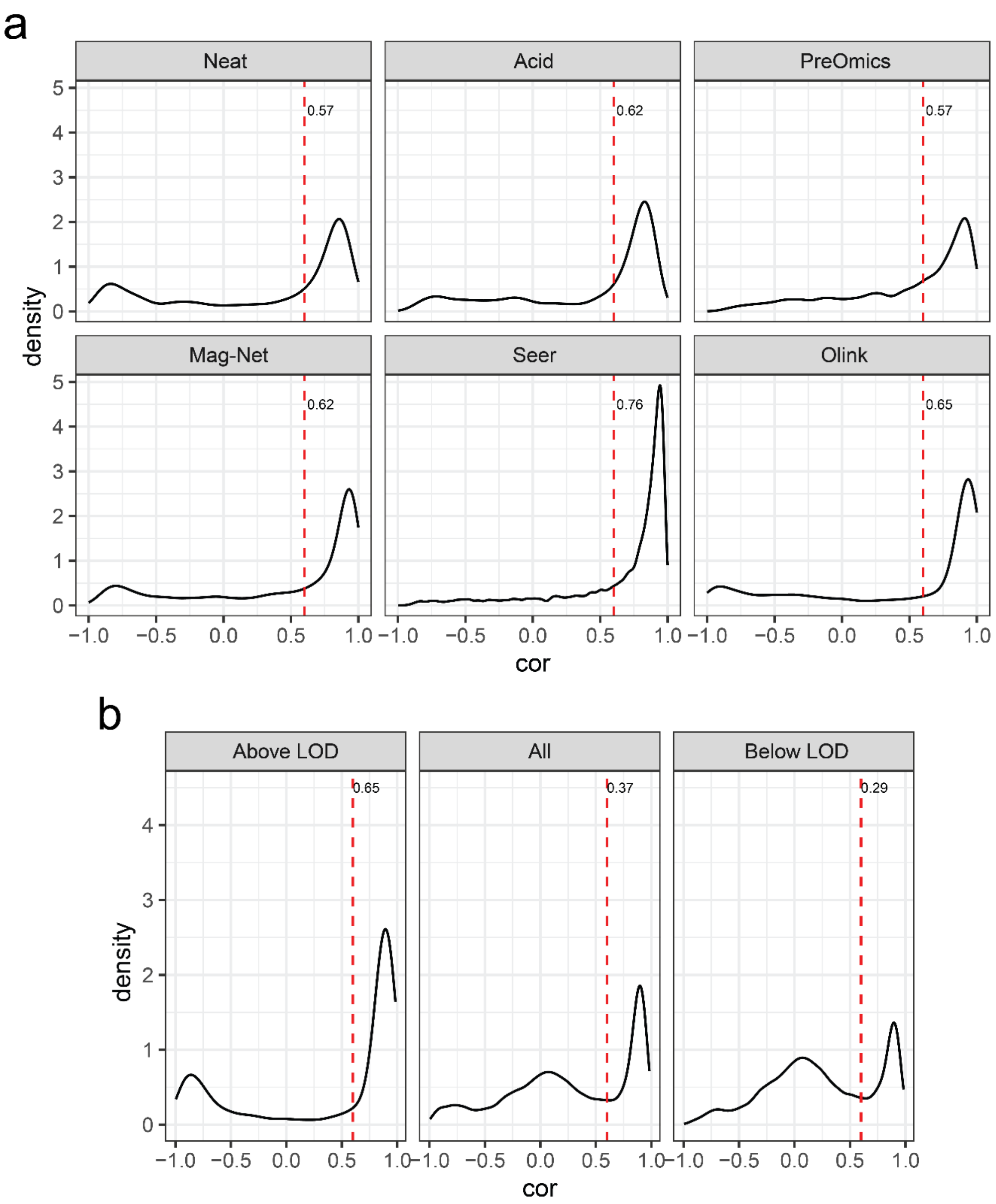


**Supplemental Figure 6 | Correlation density plots for MMCC dilutions.** (a) Correlations were calculated between the 10 triplicate human plasma dilution points and the quantity of each protein for all methods. The density of the correlations were plotted and shown is the proportion of proteins with a correlation > 0.6 to the dilution curve. For Olink the LOD cutoff was applied. (b) Correlation density plots for Olink proteins above LOD, all proteins, or only those below LOD.

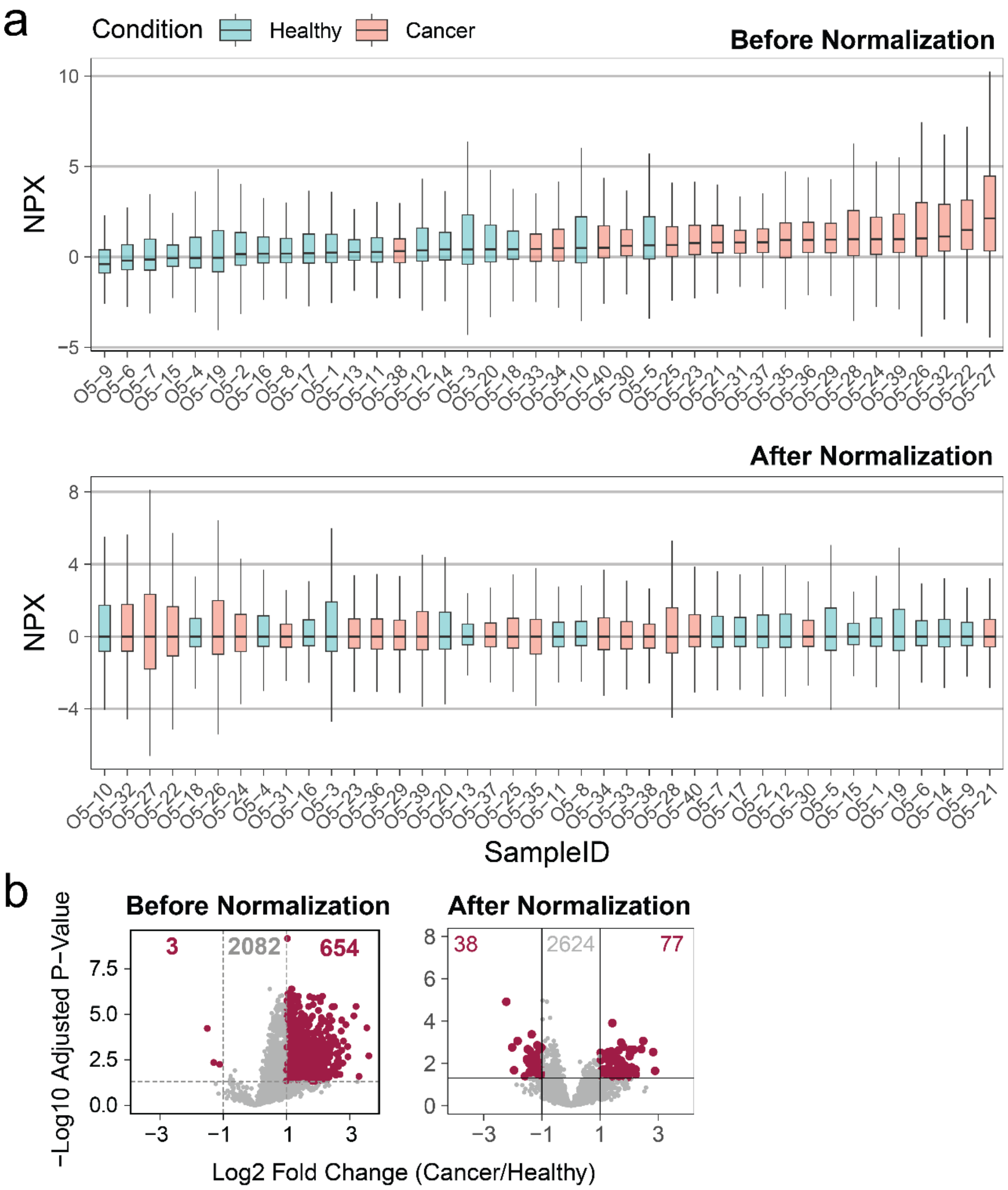


**Supplemental Figure 7 | Olink Median Normalization.** (a) For the 40-sample cancer cohort after LOD filtering and imputation, Olink data was normalized by subtracting the median NPX value for each sample from all the protein NPX values for that sample. (b) The differential expression results (from Figure 7) between Cancer and Healthy groups are no longer skewed toward up-regulation.

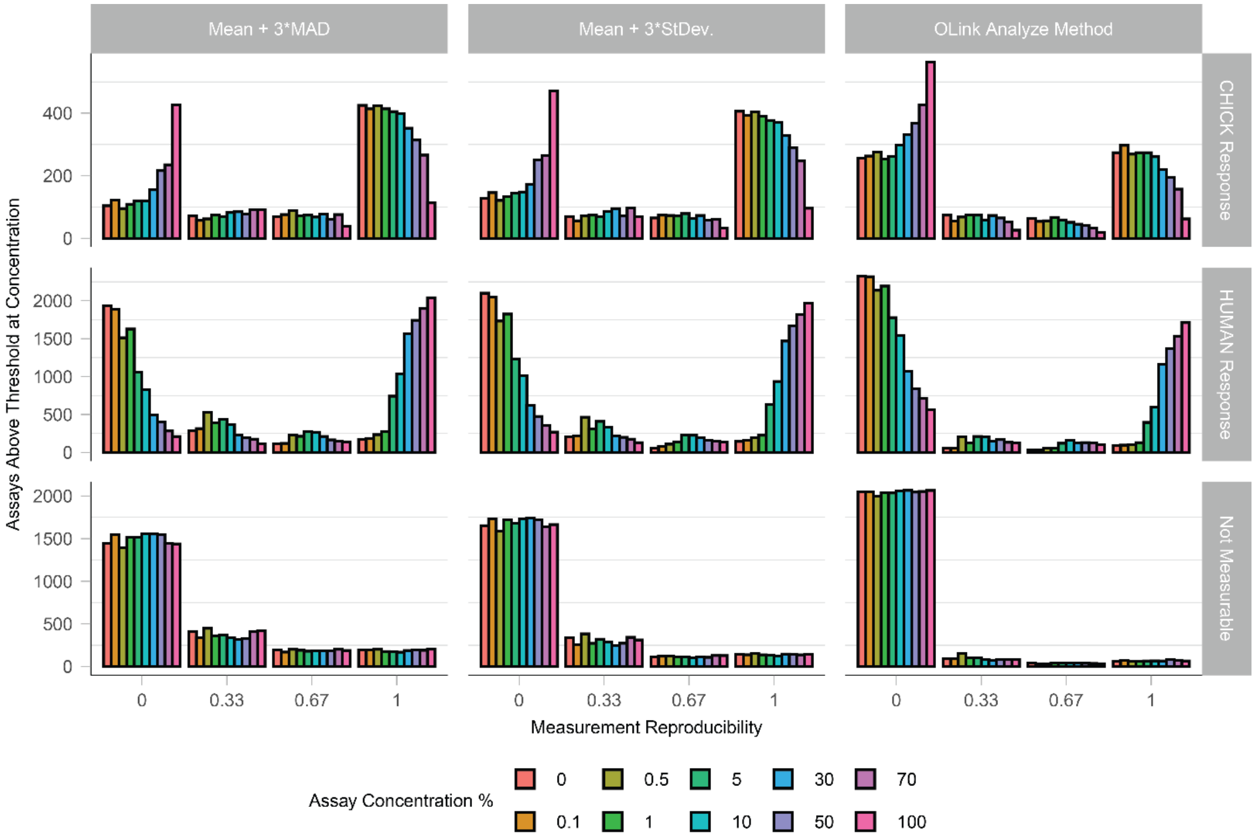


**Supplemental Figure 8 | Grouping of Olink assays according to three LOD cutoffs.** Using the negative controls built into Olink Explore HT, three signal/noise cutoffs were calculated. The rows of response were based on the calculation of LOD/LOQ in the decreasing human plasma direction or decreasing chicken plasma direction. Assays in both were considered “Not Measurable”. Within each facet, assays above the signal/noise threshold were plotted for each concentration of human plasma at each level of reproducibility (no replicates, one replicate, two replicates, or three replicates).
